# A cyclic adenosine monophosphate (cAMP)-specific phosphodiesterase interacts with begomovirus capsid proteins and modulates virus retention within its vector

**DOI:** 10.1101/2023.08.08.552480

**Authors:** Saptarshi Ghosh, Banani Mondal, Ola Jassar, Murad Ghanim, Saurabh Gautam, Rajagopalbabu Srinivasan

## Abstract

Begomoviruses are whitefly-transmitted ss-DNA viruses infecting dicotyledonous plants and contribute to major economic losses to global crop production. Invasion and establishment of an aggressive species of *B. tabaci*, known as the B cryptic species, has severely constrained vegetable production in the southeastern and southwestern United States. Disruption of genes/pathways critical for whitefly mediated transmission can be effective for the management of begomoviruses. In this study, yeast two hybrid (Y2H)-based screening of *B. tabaci* cDNA library identified a cyclic adenosine monophosphate (cAMP)-specific phosphodiesterase (PDE4) of the whitefly as an interacting partner with capsid proteins (CPs) of old- and new-world begomoviruses. Interactions of PDE4 with begomovirus CPs were validated by GST-pull-down assays, co-immunoprecipitation, and co-immunolocalization in whitefly midgut. The PDE4 family of enzymes hydrolyze cAMP and regulate intracellular cAMP levels. This study revealed that elevation of cAMP within whitefly by chemically inhibiting or gene (PDE4) silencing resulted in increased retention and transmission of begomoviruses. Similarly, decreased cAMP levels resulted in reduced begomovirus retention. The results of this study demonstrate that whitefly mediated transmission of begomoviruses is regulated by intracellular cAMP by unknown mechanisms.

**Importance:** Begomoviruses, transmitted by the sweetpotato whitefly, (*Bemisa tabaci Gennadius*), are the causal agents of many economically important plant virus diseases. Lack of host plant resistance against begomoviruses, high whitefly abundance, and whitefly’s ability to develop insecticide resistance rapidly often renders commonly used management practice ineffective. This study demonstrates how begomovirus retention within whitefly and its transmission can be modulated by altering cAMP expression of its insect vector. Naturally occurring bio-pesticides that target insect cAMPs are known. Our findings can lead to alternative strategies for the management of begomoviruses by targeting whitefly cAMP using chemicals, botanicals, or RNAi-based insecticides.

## Introduction

The sweetpotato whitefly, *Bemisia tabaci* Gennadius, is a cryptic species complex (1). *B. tabaci* is infamous for its ability to transmit several economically important plant viruses that challenge food security in the tropics and semi-tropics (2,3). Among the diverse group of viruses transmitted by *B. tabaci*, ss-DNA viruses with geminate particles belonging to the genus *Begomovirus* (family *Geminiviridae*) constitute the largest group of viruses (2). These viruses are transmitted by *B. tabaci* in a circulative and non-propagative mode. The begomovirus genomes exist in either monopartite form with a single circular ss-DNA (2.8 kb) known as DNA A or bipartite form with an additional single circular ss-DNA (∼2.8 kb) known as DNA-B (4). AV1 gene in DNA-A encodes the highly conserved capsid protein (CP), which is recognized specifically by cell surface receptors in whitefly midgut and salivary gland tissues resulting in stringent virus-vector relationships (2). Begomoviruses while circulating within the whitefly interact with multiple intracellular proteins and bacterial endosymbionts through CP, which facilitates tropism across the tissue barriers such as the midgut, salivary glands, and the hemolymph (2).

Begomoviruses are broadly classified as new-world or old-world based on their origin and distinguished by the absence of AV2 gene in the DNA A component of the former. Increased agricultural trade between the new- and old-world in the past decades has led to exchange and establishment of exotic begomoviruses and invasive whitefly cryptic species in non-native geographical areas (5). A highly invasive cryptic species of *B. tabaci,* B cryptic species (also known as Middle East Asia Minor MEAM1), invaded the US in the late ’80s and established itself across the southeastern and southwestern states, and displaced the indigenous whitefly cryptic species ‘A’ (6–8). In these regions within the United States, B cryptic species of *B. tabaci* constrains vegetable production by efficiently transmitting new-world begomoviruses such as cucurbit leaf crumple virus (CuLCrV) that infects cucurbit crops and sida golden mosaic virus (SiGMV) that infects snap bean (9,10). The B cryptic species also efficiently transmits old-world viruses such as tomato yellow leaf curl virus (TYLCV) that infects tomato (5,11,12). Management of begomoviruses is challenging due to the limited availability of host plant resistance (13), consequently, there is a heavy reliance on cultural and chemical practices (14,15). However, extremely high whitefly abundance during some seasons (for instance, fall season in the southeastern US) renders insecticidal management of virus transmission ineffective (15–18). There is a dire need for novel and sustainable strategies for the management of whitefly-transmitted begomoviruses in vegetables. Identification of whitefly proteins/pathways, involved in the transmission of begomoviruses, is important for the conception of novel virus management strategies. Although multiple whitefly proteins with inhibitory roles (19–23) against begomoviruses have been identified, an exact mechanism or pathway that can be targeted to reduce virus transmission remains unknown.

This study used yeast-two-hybrid (Y2H) based screening of *B. tabaci* (B biotype) cDNA library with the CP of CuLCrV as bait and identified cAMP specific 3’, 5’-cyclic nucleotide phosphodiesterase of the whitefly as an interacting partner. The cAMP-specific phosphodiesterases hydrolyze cAMP –an intracellular second messenger of extracellular ligand action (24), and play an important role in modulating inflammatory responses (25). In this study, cAMP levels were elevated and diminished within the whitefly using chemical inhibitors and gene-silencing. This study also demonstrated that the retention and transmission of begomoviruses are directly dependent on intracellular cAMP levels. These outcomes have important implications for the management of epidemics of begomoviruses.

## Materials and methods

### Maintenance of *B. tabaci* B cryptic species population and virus isolates

*B. tabaci* (B cryptic species) was maintained on cotton plants inside insect proof cages at 25□6 □C, 60% RH, and 14L:10D (18). CuLCrV, TYLCV and SiGMV virus isolates were maintained using whitefly mediated inoculation in squash (cv. F1 Goldstar hybrid), tomato (cv. Florida 47) and Sida, respectively (10,16,18).

### Construction of *B. tabaci* B cryptic species cDNA library and Y2H screening

Total RNA (2□g) extracted from *B. tabaci* adults was used for the synthesis of first strand cDNA utilizing the make your own mate & plate library system with CDS III adaptor primer and smart III oligo (template switching) as per manufacturer’s instructions (Takara Bio, San Jose, CA, USA). The first strand cDNA was used as template for long distance PCR with 5’ and 3’ primers, purified using chroma spin + TE 400 columns, and co-transformed into competent Y187 yeast cells for gal4 AD-*B. tabaci* B cDNA library construction as previously described (26). The pGBKT7 construct with CP of CuLCrV as insert (5) was transformed in Y2H Gold strain of yeast and screened against the *B. tabaci* B library on (–ALTH) minimal media (5). Prey plasmids from colonies developing blue color on X-alpha-gal supplemented – ALTH media were recovered and sequenced from both directions by Sanger sequencing. Nucleotide sequences were translated *in silico* in frame with the GAL4 activation domain and annotated using BLASTp, CDD, and Pfam databases. PDE4 gene starting from ATG was PCR amplified using specific primers (Table 1) from cDNA of MEAM1 and cloned into pGADT7 and was screened against the CPs of new-world viruses (CuLCrV, SiGMV), old -world virus (TYLCV) and empty pGBKT7 by one-to-one mating assays (5) on -ALTH media and lesser stringent -LTH minimal media, respectively.

**Primer table 1.**
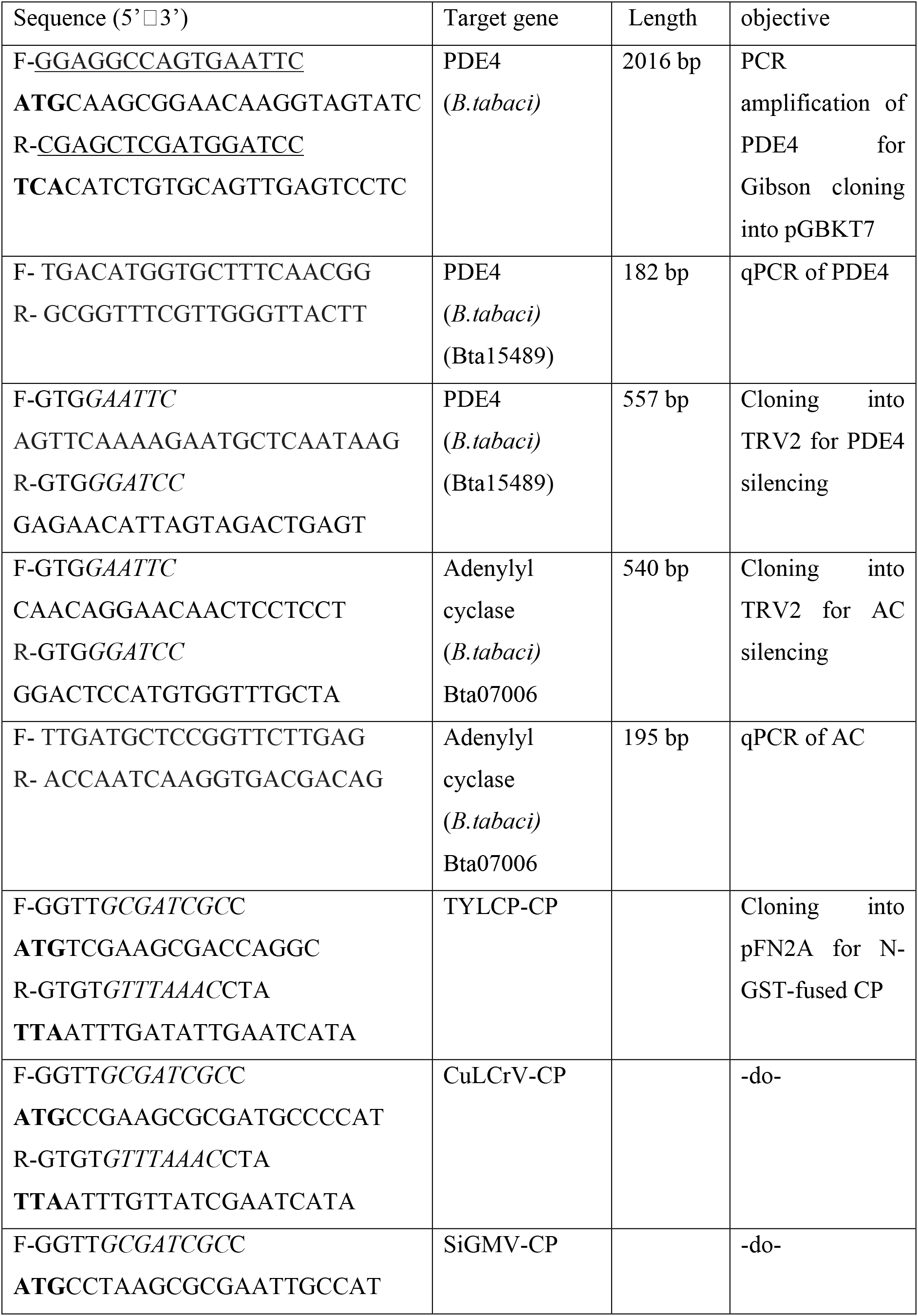
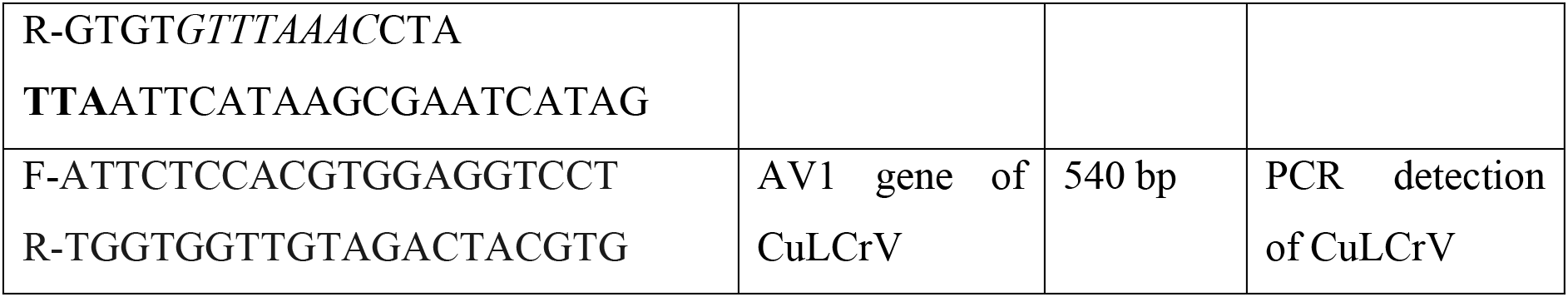

### Pull-down assay of PDE4 using glutathione-S-transferase (GST) tagged CPs of CuLCrV, SiGMV, and TYLCV

CP genes of CuLCrV, SiGMV and TYLCV were cloned in pFN2A vector (Promega) using *Sgf*I and *Pme*I restriction enzymes to express them as fused proteins with N-terminal GST tag. PDE4 gene was inserted in pRSET A (ThermoFisher scientific, USA) using the *Bam*HI and *Eco*RI restriction sites to express it as fused proteins with N-terminal 6X-histidine tag. All the constructs were transformed to BL21 DE3 pLysS strain and protein expression was induced with 0.4 mM IPTG at 28□C for 5 hours. Bacterial lysates expressing fused GST-CuLCrV / SiGMV / TYLCV CP proteins and GST control were incubated with MagneGST□ particles (cat#V8600, MagneGST protein purification system, Promega, Madison, WI, USA) to immobilize the GST tagged proteins as per manufacturer’s instructions. MagneGST particles immobilized with the respective GST tagged proteins were incubated with bacterial lysates expressing fused 6xhis-PDE4 overnight at 4□C, washed three times with 400□l of binding/wash buffer followed by boiling with 35□l of 1x Laemmli buffer. The proteins were electrophoresed on 12% polyacrylamide gels, transferred to nitrocellulose membranes, the membrane was cut into two pieces between 55 and 70 kD ladder bands followed by overnight incubation at 4□C with either monoclonal anti-histidine (1:2000) (cat#MAB3834, Millipore Sigma, USA) or anti-GST (1:1000) (cat#8-326, Thermofisher scientific) antibodies. Histidine tagged PDE4 and GST tagged CP proteins were detected chromogenically using anti-mouse secondary antibodies labelled with alkaline phosphatase (1:30000) (cat#A3562, Millipore Sigma, Burlington, MA, USA) and western blue stabilized substrate for alkaline phosphatase (Promega, Madison, WI, USA).

### Coimmunoprecipitation of the whitefly PDE4 protein using polyclonal anti-CP antibody

Viruliferous whitefly adults (∼200) reared on SiGMV or CuLCrV infected sida or cucurbit plants were homogenized in 600 □l of CytoBuster protein extraction reagent (Millipore Sigma, Burlington, MA, USA) with 1X protease inhibitor cocktail (Cat#G6521, Promega, USA) and incubated at 4□C with gentle shaking for 1.5 hours. The samples were centrifuged at 10000 rpm for 10 minutes at 4□C, the supernatant was collected, protein quantities were assessed by Bradford assay, and were stored in aliquots at −80□C. Protein A magnetic beads (Cat#S1425S, New England Biolabs Inc, Ipswich, MA, USA) were aliquoted (20 □l) into 1.5 ml microcentrifuge tubes, washed twice with 50 □l PBS (pH 7.4), and incubated with 1:100 dilutions of either anti SiGMV-CP polyclonal antibody pre-immune antiserum in PBS with 1% BSA at 4□C for 2 hours with gentle shaking. The beads were washed thrice with PBST (PBS+0.05% tween 20) and incubated with 70 ng of extracted whitefly protein overnight at 4□C with gentle shaking. The beads were washed thrice with PBST, bound proteins were eluted by boiling with 25 □l of 3X-Laemmli buffer and electrophoresed on 10% polyacrylamide gel. PDE4 protein co-immunoprecipitated by the anti-CP antibody was detected by western blot using 1:5000 dilution of a polyclonal anti-PDE4 antibody generated by immunizing two rabbits with a 243 amino acid long partial PDE4 protein (NDESVLENHHLVVEFKLLQKEGCDIFINLSKKQKQTLRKMVIDMVLSTDMSKHMSLLADLKAMVETKKVAGSGVLLLDNYTDRIQVLENLVHCADLSNPTKPLPLYRRWVDLLMEEFFQQGDKEREQNLDISPMCDRHSATIEKSQVGFIDYIVHPLWETWADLVHPDAQEILDMLEENRDWYQSMIPPSPPVNEGENRLDSDVEEGEESEPPNPNPPPVPQDSSIRFQVTLEEGDEDSTAQM) expressed in *E. coli* by (CUSABIO, Houston, Texas, USA). A goat anti rabbit secondary antibody (1:30000) conjugated to alkaline phosphatase (Cat#A3687, Millipore Sigma, Burlington, MA, USA) and western blue stabilized substrate for alkaline phosphatase (Promega, Madison, WI, USA) was used for chromogenic visualization of protein bands.

### Immunolocalization of PDE4 and TYLCV in *B. tabaci* midgut

Viruliferous whitefly reared on TYLCV-infected plants were dissected in PBS and the midguts were fixed in 4% paraformaldehyde for 30 minutes at room temperature followed by permeabilization with 0.1% Triton X-100 for 30 minutes at room temperature. The midgut samples were washed with PBST thrice, incubated for one hour in blocking buffer (PBST+0.5% BSA) followed by overnight incubation with anti TYLCV-CP polyclonal antibody (1:500) at 4□C. The midgut samples were washed thrice with PBST and incubated for two hours at room temperature with cy5 conjugated donkey anti-rabbit secondary antibody (Jackson ImmunoResearch Laboratories, PA, USA) diluted (1:1000) in blocking buffer. The midgut samples were then washed thrice with PBST and incubated overnight at 4□C in blocking buffer containing anti-PDE4 polyclonal antibody (1:1000). Subsequently, the midgut samples were washed thrice with PBST and incubated for two hours at room temperature with cy3 conjugated donkey anti-rabbit secondary antibody (Jackson Laboratories) diluted (1:1000) in blocking buffer. The midgut samples incubated with cy3/cy5 conjugated secondary antibody without exposure to the anti-CP or anti-PDE4 antibody were used as negative control. Finally, the midgut samples were washed thrice with PBST and mounted with a buffer (PBS+10 ng DAPI) on a glass slide with a cover slip, sealed using nail polish and viewed under an Olympus IX81 laser scanning confocal microscope.

### Inhibition of cAMP phosphodiesterase (PDE4) and quantitation of cellular cAMP in whitefly

Rolipram (cat#0905, Tocris Bioscience, Bristol, UK), a selective inhibitor of cAMP phosphodiesterase, was used to inhibit PDE4 and stimulate cellular cAMP within whitefly adults. The increase in cellular cAMP post feeding with rolipram was validated by quantitation of cAMP from adult whitefly samples using the Direct cAMP ELISA kit (cat#ADI-900-066A, ENZO Life Sciences, Farmingdale, NY, USA). Whitefly adults after 48 hours of feeding on 20% sucrose diets containing either 200□M rolipram or 0.8% ethanol were collected and immediately snap-frozen in liquid nitrogen. Fifty females were homogenized in 250□l of 0.1N HCl and kept on ice. An aliquot (25□l) of this lysate was used for protein quantitation using Pierce Coomassie protein assay kit (cat#23200, ThermoFisher scientific). The remaining lysate was centrifuged at 5000xg at 4C to pellet debris. Two technical replicates, each with 100□l of the supernatant was used for estimation of cAMP in the samples alongside cAMP standards (non-acetylated) using the competitive ELISA kit as per manufacturer’s instructions. Sample cAMP concentrations (pmol/ml) obtained by fitting in the OD values in the regression equation generated by plotting the standards was normalized to the protein content (pmol/mg proteins) in the lysate. Five separate biological replicates from each treatment were estimated for cAMP and means of four biological replicates (1 outlier removed) were analyzed using students t-test.

To know if cellular cAMP varies with acquisition of begomoviruses, cAMP levels were compared between viruliferous and non-virulierous whitefly adults. Whitefly adults with 72 hours of acquisition access period on CuLCrV infected and non-infected squash plants were collected and snap frozen. Cyclic AMP was quantified from a pool of 50 female insects as described before using the cAMP specific ELISA kit.

### Effect of rolipram on virus retention by whiteflies

Whitefly adults were fed on 20% sucrose diet containing either 200 □M rolipram (solubilized in ethanol) or 0.8% ethanol as control for 48 hours. The diet fed whiteflies were allowed 24 hours of acquisition access to either CuLCrV-infected squash or TYLCV-infected tomato plantlets excised from a single infected source plant, followed by overnight gut clearing on cotton plants. Total DNA extracted from pools of 10 gut cleared whitefly adults was used as template for relative quantitation of CuLCrV and TYLCV retained using CuLCrV-qF/R and TYLCV-V2F/R and normalized to the □-tubulin gene of the whitefly as previously described (5) using the □□Ct method. Significant differences of means were compared using one-way ANOVA.

### Inhibition of adenylyl cyclase to decrease cellular cAMP and effect on virus retention

SQ22536 (cat#1435, Tocris Bioscience), an inhibitor of adenylyl cyclase (AC) was used to diminish cellular cAMP in the whitefly. Whitefly adults were fed for 48 hours on 20% sucrose diet containing 200□M of SQ22536 (solubilized in ethanol) or 0.8% ethanol as control. The diet fed whiteflies were allowed 24 hours of acquisition access to CuLCrV or TYLCV infected squash/tomato plantlets, gut cleared by overnight feeding on cotton plants followed by quantitation of CuLCrV/TYLCV retained by qPCR as described above. Significant differences of means were compared by one-way ANOVA.

### Virus induced gene silencing of PDE4, adenylyl cyclase (AC), and effect on virus retention

To further validate the role of cAMP, virus induced gene silencing using tobacco rattle virus-based vectors were used to silence PDE4 and AC genes in the whitefly as previously described (5,27). A 557 and 540 bp fragment of the PDE4 (Table1) and AC genes (Table1), respectively was cloned into the mcs region of pTRV2 vector using *Eco*RI and *Bam*HI restriction enzymes. The PDE4_TRV2, AC_pTRV2, pTRV2 control and pTRV1 were transformed separately into LBA4404 strain of *Agrobacterium tumefaciens*, induced for 3-5 hours at room temperature. pTRV1 and PDE4_pTRV2 or AC_TRV2 or TRV2 control cultures were mixed in 1:1 proportion (v/v) and infiltrated within the leaves of 12 days old tomato seedlings using a 1 ml syringe. Transformation of tomato plants was confirmed with detection of PDE4/AC on systemic leaves 12 days post inoculation by RT-PCR. *Bemisia tabaci* colonies were set up by releasing adult whiteflies on transformed plants for 10 days. Relative expression of PDE4 and AC normalized to the whitefly □-tubulin in F1 adult populations of whitefly feeding on plants inoculated with pTRV1 + PDE4/AC-pTRV2 constructs or TRV1 + TRV2 (control) constructs was quantitated by qPCR, and their means were compared by one-way ANOVA to confirm silencing. Following confirmation of silencing, F1 adults were given acquisition access to CuLCrV/TYLCV infected squash/tomato plants for 24 hours, gut cleared by overnight feeding on cotton plants, and relative amounts of virus retained was quantitated by qPCR. Significant differences of means were compared by one-way ANOVA.

### Effect of rolipram/SQ22536 on transmission of CuLCrV by *B. tabaci*

*Bemisia tabaci* adults (2-7 days old) reared on cotton plants were collected and provided with a 48-hour feeding access on 20% sucrose diet containing 200 □M of rolipram or 0.8% ethanol as control. Whitefly adults were then released separately onto a squash leaf excised from a single CuLCrV infected squash plant for an acquisition access period of 48 hours. Adults collected after acquisition access from the infected leaf were allowed life-long inoculation access (10 adults/plant) onto young squash seedlings (cv. F1 Goldstar hybrid1-2 true leaf stage) in three separate replicated experiments. Total DNA was extracted from inoculated squash plants 15 days post inoculation and PCR indexed for CuLCrV infection using primers targeting the CP (AV1) gene (F-ATTCTCCACGTGGAGGTCCT, R-TGGTGGTTGTAGACTACGTG, 540 bp). Similarly, whitefly adults fed for 48 hours on 20% sucrose diet with and without 200 □M SQ22536 were provided with a 48-hour acquisition access period and an inoculation access on non-infected cucurbit plants in two replicates containing 12 plants each. The plants were tested for CuLCrV 15 days post inoculation by PCR. Significant differences in transmission efficiency were inferred by analysis using Chi-square test.

Plant samples confirmed infected with CuLCrV in the transmission assays were further used for absolute quantitation of CuLCrV using CuLCrV-QF/R primers and CuLCrV-hydrolysis probe (11). The means of virus copy numbers/ng of DNA template between rolipram/SQ22536 and control plants were analyzed using Wilcoxon-rank sum test.

## Results

### Identification of PDE4 as an interacting partner with CPs of begomoviruses

Y2H screening of MEAM1 cDNA library with CP of CuLCrV as bait resulted in identification of diploid yeast colonies developing on -ALTH minimal media. Inserts within colonies included a 2065 bp CDS (Fig. 1, OR396905). The CDS encoded a protein with a phosphodiesterase 4 N-terminal conserved region (PDE4_NCR) domain and a cAMP specific 3’, 5’-cyclic nucleotide phosphodiesterase (PDEase_I) domain. BLAST analysis of the nucleotide sequence revealed >99.5% similarity with cAMP specific 3’ 5’-cyclic nucleotide phosphodiesterase of *B. tabaci* (<u>XM_019060127.1</u>, <u>XM_019060128.1</u>, <u>XM_019060129.1</u>). *Bemisia tabaci* phosphodiesterase transcripts identified in this study will be referred as PDE4 here onwards. BLASTP (NCBI/FlyBase) analysis of translated amino acid sequence showed high identity with cAMP-specific 3’, 5’-cyclic phosphodiesterase of diverse insects. The dunce (dnc) gene of *Drosophila melanogaster* encoding cAMP-specific 3’,5’-cyclic phosphodiesterase, isoform I (phosphodiesterase 4 subfamily), shared > 74% amino acid sequence identity with the *B. tabaci* homolog. The *B. tabaci* PDE4 transcript identified in this study had additional 5’ sequences (Fig. 1), which were missing in the *B. tabaci* sequences available on Genbank, indicating that the Genbank sequences were 5’ partial.

**Figure 1:**
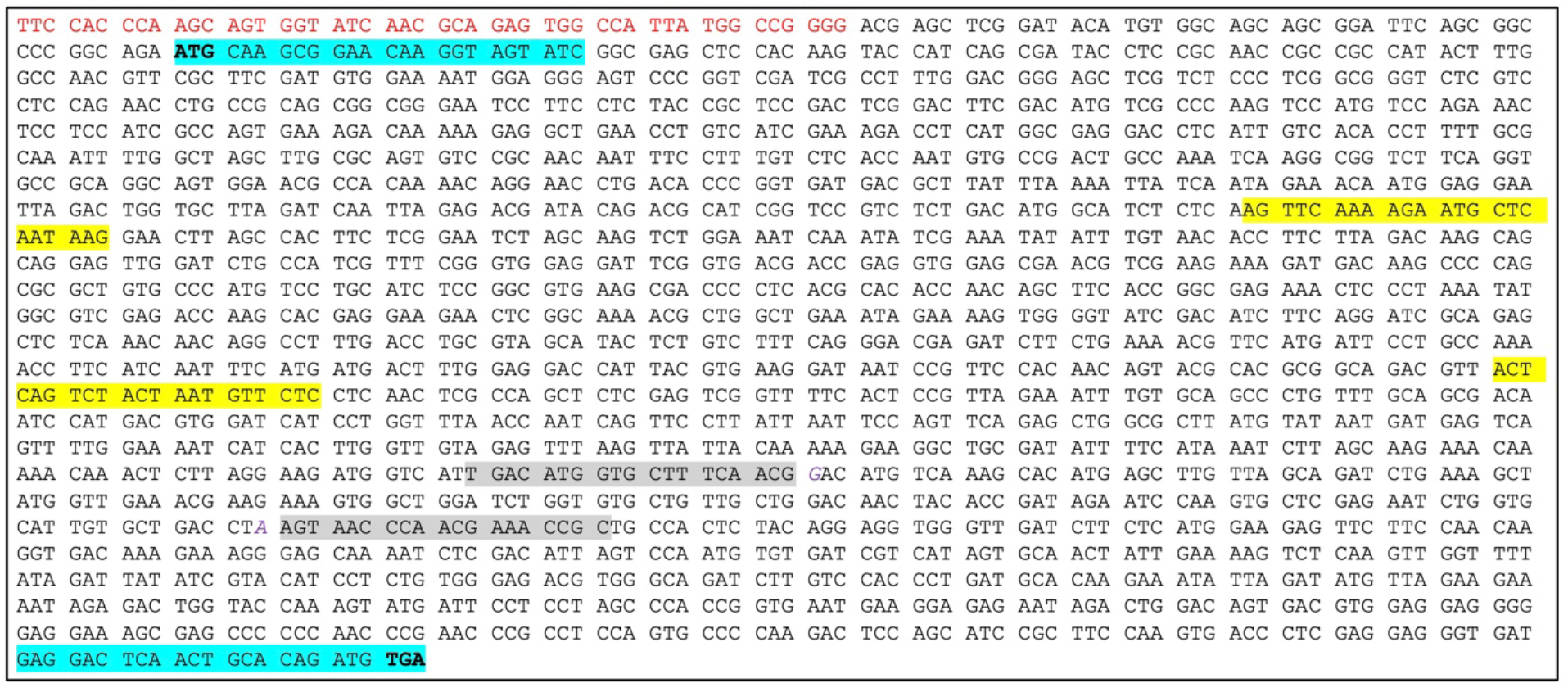
Nucleotide sequence of the PDE4 construct identified by Y2H screening of *B. tabaci* (B cryptic species) cDNA library against CP of CuLCrV. Nucleotide sequences of the smart III oligo is denoted in red, primers used for PCR amplification of PDE4 (2016 bp) for Y2H mating/pull-down assay, qPCR (182 bp) and dsRNA (557 bp) TRV2-construct are highlighted in blue, grey, and yellow, respectively. Bold letters denote start and stop codons of the PDE4 gene used in this study.

### Validation of interaction of PDE4 with CPs of begomoviruses

#### Y2H protein interactions of PDE4 with CPs of begomoviruses

A 2016 bp nucleotide (Fig.1) fragment (from start to stop codon) encoding a 671 amino acid long PDE4 protein was PCR amplified using *B. tabaci* cDNA as template and cloned in pGADT7 to encode a fused protein with N-terminal gal4 activation domain (AD). The first 15 amino acid sequences (MQAEQGSIGELHKYH) of the translated 671 amino acids long PDE4 protein of *B. tabaci* matched with the initial sequences of complete cAMP specific 3’, 5’-cyclic nucleotide phosphodiesterase proteins of aphids (<u>XP_026814909.1</u>, <u>XP_027852822.2</u>), thrips (<u>XP_034240939.1</u>), wasps (<u>XP_044579040.1</u>, <u>XP_034946957.1</u>) and beetles (<u>XP_008198075.1</u>, <u>XP_018320283.1</u>) indicating that a full length gene of PDE4 was used in this study. The gal4 AD-PDE4 construct was screened against gal4 BD-CP (CuLCrV/SiGMV/TYLCV) or empty pGBKT7 by one-to-one Y2H mating analysis. Diploid yeast colonies containing gal4 AD-PDE4 and gal4 BD-CP constructs of the tested new-world viruses (CuLCrV and SiGMV) turned blue (Fig. 2A) on stringent minimal media (-ALTH + X-□gal), whereas colonies with TYLCV CP were observed (Fig. 2B) only on the less stringent (-LTH) medium. No growth was observed for yeast containing PDE4 and empty pGBKT7 constructs in either medium (Fig. 2 A, B). These results indicate that PDE4 protein of *B. tabaci* interacts with the CPs of begomoviruses.

**Figure 2:**
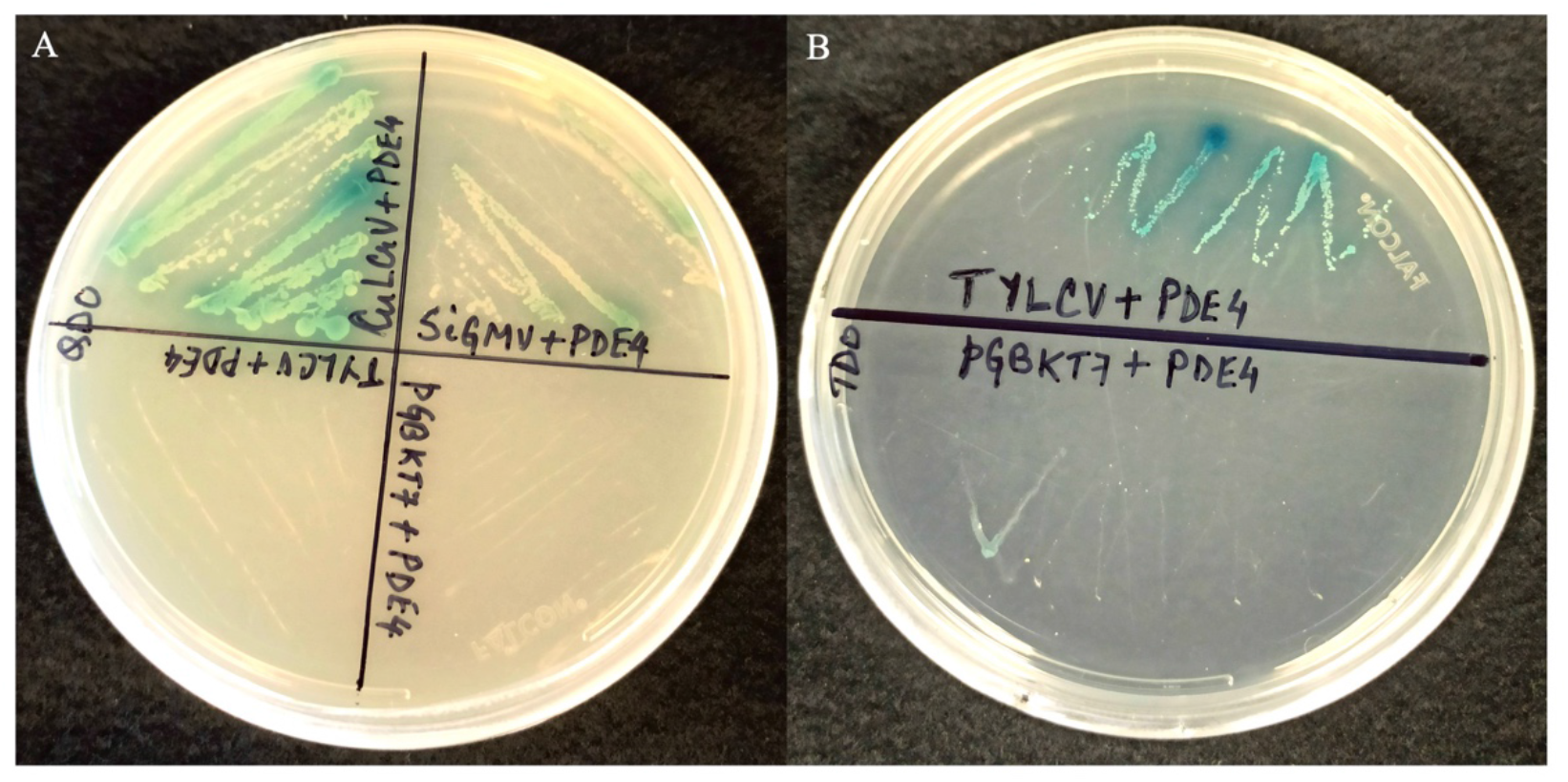
Interaction of PDE4 with CP proteins of CuLCrV, SiGMV, TYLCV and empty pGBKT7 by one-one Y2H mating assays on (A) stringent -ALTH and (B) less stringent -LTH minimal media supplemented with α-Xgal.

#### Pull-down assay of PDE4 using CPs of begomoviruses

Interactions between the begomovirus CPs and PDE4 protein was validated by GST pull-down assay using CPs of CuLCrV, SiGMV, and TYLCV expressed as fused proteins (baits) with N-terminal GST to capture expressed 6x-his-PDE4 proteins from soluble fractions of the bacterial lysate. Results showed that GST fused CPs of CuLCrV, SiGMV, and TYLCV bound to PDE4 of *B. tabaci,* whereas no binding was detected between GST and PDE4 (Fig. 3A).

**Figure 3:**
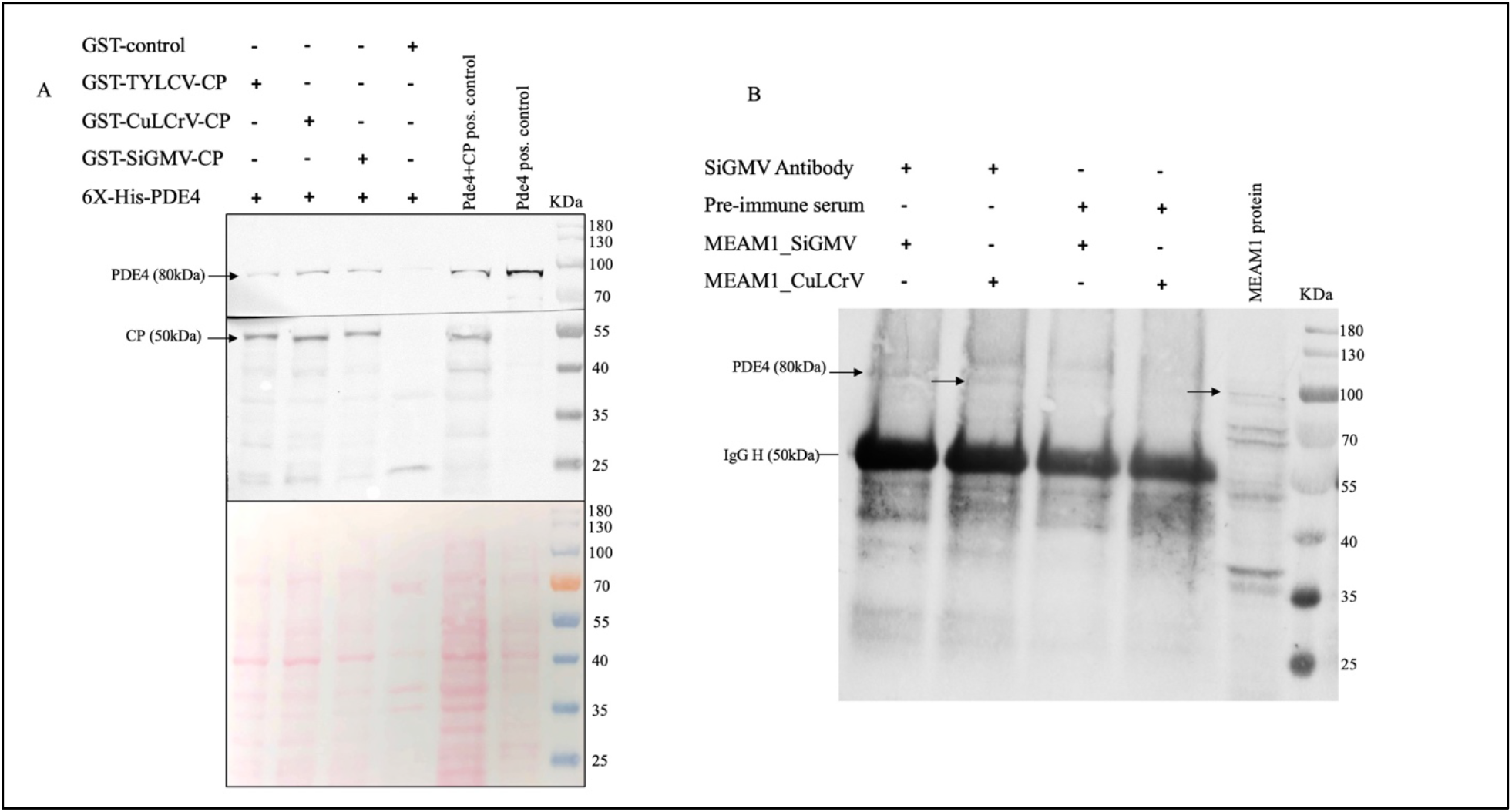
Validation of interactions between begomovirus CPs and PDE4. A) Pull-down assay of *B. tabaci* PDE4 protein from bacterial lysates expressing 6X-his-PDE4 using GST fused CPs of TYLCV, CuLCrV and SiGMV as baits. (A) Mixture of bacterial lysates expressing the CP and PDE4 and only PDE4 was used as positive control for detection. The ponceau stained proteins show the protein amounts of the individual lanes. (B) Co-immunoprecipitation of PDE4 protein from soluble fraction of viruliferous (CuLCrV/SiGMV) MEAM1 proteins using polyclonal anti-SiGMV CP antibody or pre-immune serum as control.

#### Co-immunoprecipitation of PDE4

PDE4 proteins were captured (Fig. 3B) from total soluble proteins of viruliferous whitefly adults (SiGMV/CuLCrV) using polyclonal anti-SiGMV-CP antibody but not with the pre-immune serum.

#### PDE4 -begomovirus CP interaction in the whitefly midgut

PDE4 proteins were detected in the midgut (Fig. 4) of viruliferous (TYLCV) *B. tabaci* adults using polyclonal anti-PDE4 antibody and co-localized with TYLCV. The co-localization appeared in the cellular cytoplasm around the nuclei (60X magnification), while in the negative control using only secondary antibodies did not show any localization of either protein.

**Figure 4:**
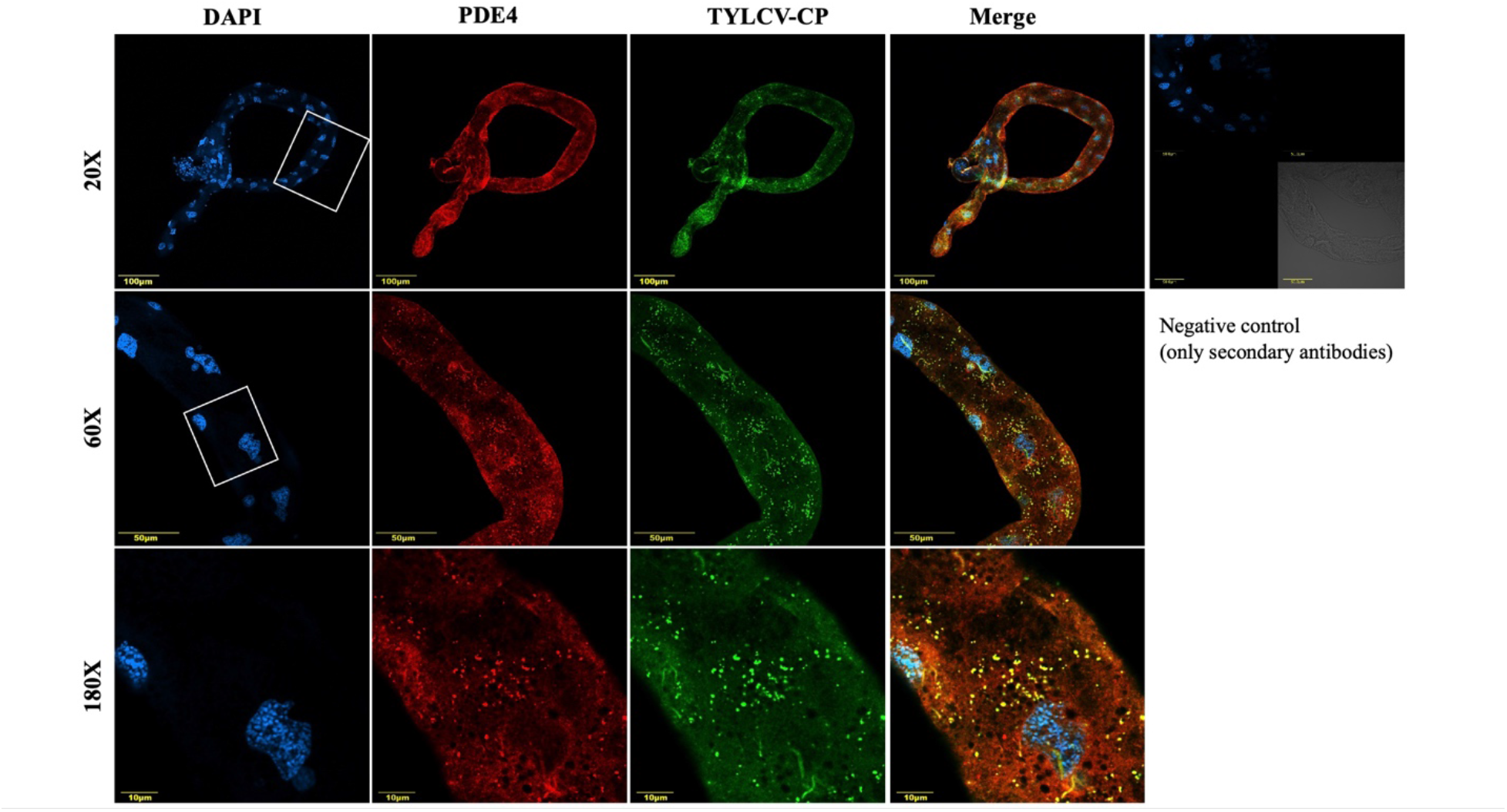
Localization of PDE4 and TYLCV in the midgut of *B. tabaci* reared on TYLCV-infected tomato plants. TYLCV was detected using polyclonal anti-TYLCV CP antibody and cy5 (green) conjugated secondary antibody. PDE4 was detected using polyclonal anti-PDE4 antibody and cy3 (red) conjugated secondary antibody. The overlay of green and red channels is indicated in yellow and midgut cell nuclei, stained with DAPI, are indicated in blue. The gut area magnified to 60X and 180X are depicted in white square boxes. Midguts incubated with cy3/cy5 conjugated secondary antibody without the primary antibody were used as negative control.

### Retention and transmission of begomoviruses are directly affected by cAMP levels within *B. tabaci*

#### Inhibition of PDE4 by rolipram

Sucrose diet supplemented with rolipram (200□M) was fed to whitefly adults for 48 hours to inhibit PDE4 and elevate intracellular cAMP. Mean cAMP concentration quantified by competitive immunoassay was significantly higher (*P*=0.038, *t*=2.65, df=6) in *B. tabaci* adults feeding on rolipram supplemented diet compared with adults on the control diet (Fig. 5A).

**Figure 5:**
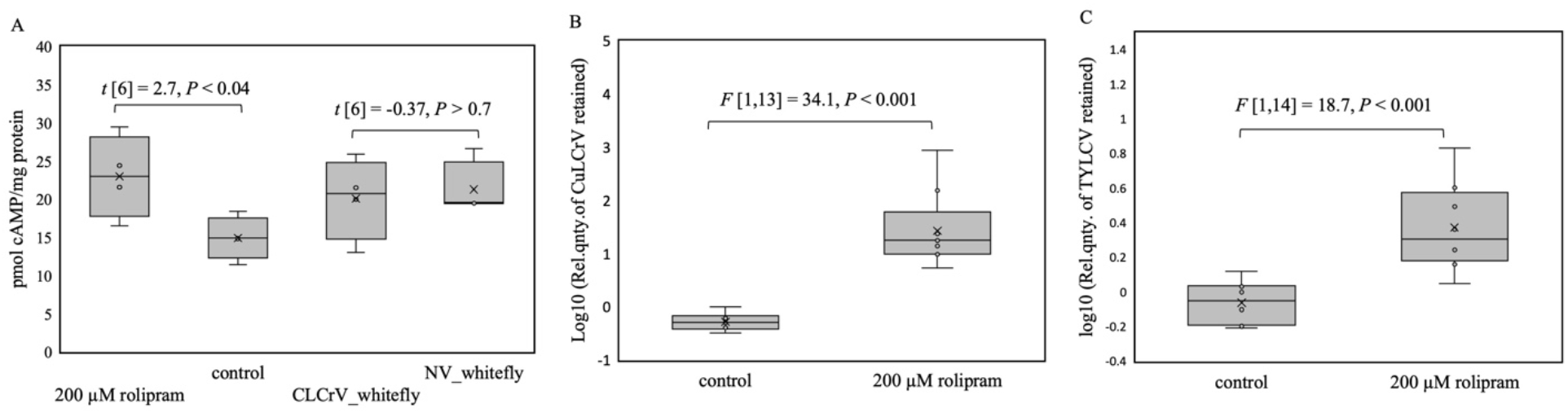
(A) Quantitation of cAMP (pico moles/mg whitefly protein) in *B. tabaci* adults either fed for 48 hours on 20% sucrose diet with/without 200 □M rolipram or whitefly adults with 72 hours of acquisition access to CuLCrV infected (CLCrV_whitefly) or non-infected squash (NV_whitefly) plants. (B) Relative quantities of CuLCrV and (C) TYLCV retained (normalized to the □-tubulin gene of the whitefly) in whitefly adults fed for 48 hours on 20% sucrose diet (control) or 20% sucrose diet supplemented with 200 □M rolipram after 24 hours of acquisition access to infected squash/tomato plants and overnight gut clearing on cotton plants.

#### Cyclic AMP levels between viruliferous and non-viruliferous whitefly adults were unaltered

Comparison of cAMP levels by competitive immunoassay between viruliferous and non-viruliferous whitefly adults 72 hours post acquisition access to CuLCrV infected plants showed no significant difference in cAMP levels upon virus acquisition (Fig. 5A).

#### Modulation of cAMP by feeding rolipram or SQ22536 alters begomovirus retention in whitefly adults

The effect of increased cAMP on begomovirus retention was investigated by comparing virus retention in whitefly adults provided with an acquisition access period (48 hours) on either rolipram (200□M) or control diet and allowed an acquisition access period (24 hours) on CuLCrV/TYLCV infected plants. Whitefly adults that fed on rolipram diet prior to acquisition access retained significantly higher quantities of CuLCrV (Fig. 5B) and TYLCV (Fig. 5C) compared with adults that fed on control diet.

Whitefly adults that fed on diets containing SQ22536 (200□M), an inhibitor of adenylate cyclase (AC), had reduced cAMP levels. When whitefly samples, feeding on SQ22536 for 48 hours, were provided with an acquisition access on virus-infected plants, they retained reduced quantities of CuLCrV (Fig. 6A) and TYLCV (Fig. 6B) compared with adults on the control diet.

**Figure 6:**
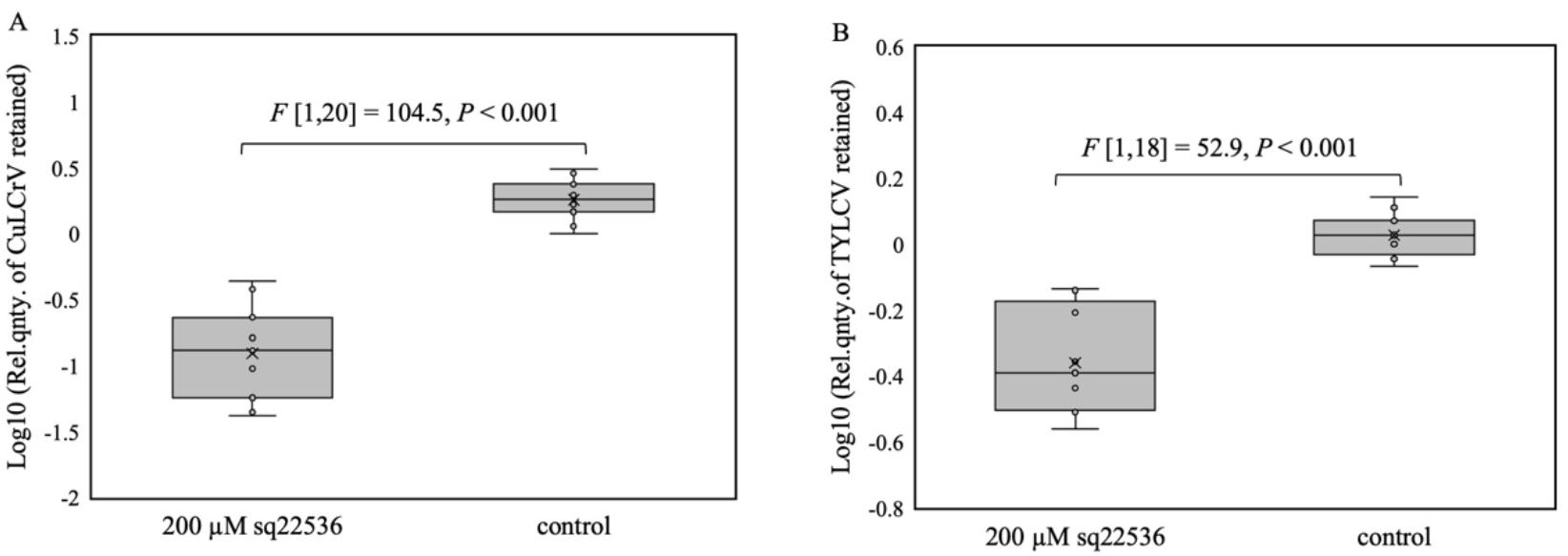
Relative quantities of (A) CuLCrV and (B) TYLCV retained (normalized to the □-tubulin gene of the whitefly) in *B. tabaci* adults fed for 48 hours on 20% sucrose diet (control) or 20% sucrose diet supplemented with 200 □M SQ22536 after 24 hours of acquisition access to infected squash/tomato plants and overnight gut clearing on cotton plants.

#### Gene silencing of PDE4 and AC alters begomovirus retention

Recombinant tobacco rattle virus (TRV) vectors were used for silencing PDE4 (Fig. 7A) and AC (Fig.7B) genes of F1 whitefly adults. *PDE4* silenced F1 adults (PDE4_TRV2) retained higher quantities of CuLCrV (Fig. 8A) and TYLCV (Fig. 8B) compared with control (TRV2) 24 hours post acquisition access on infected plants and overnight gut clearing. In contrast, AC silenced (AC_TRV2) F1 adults retained reduced amounts of CuLCrV (Fig. 8C) than control (TRV2).

**Figure 7:**
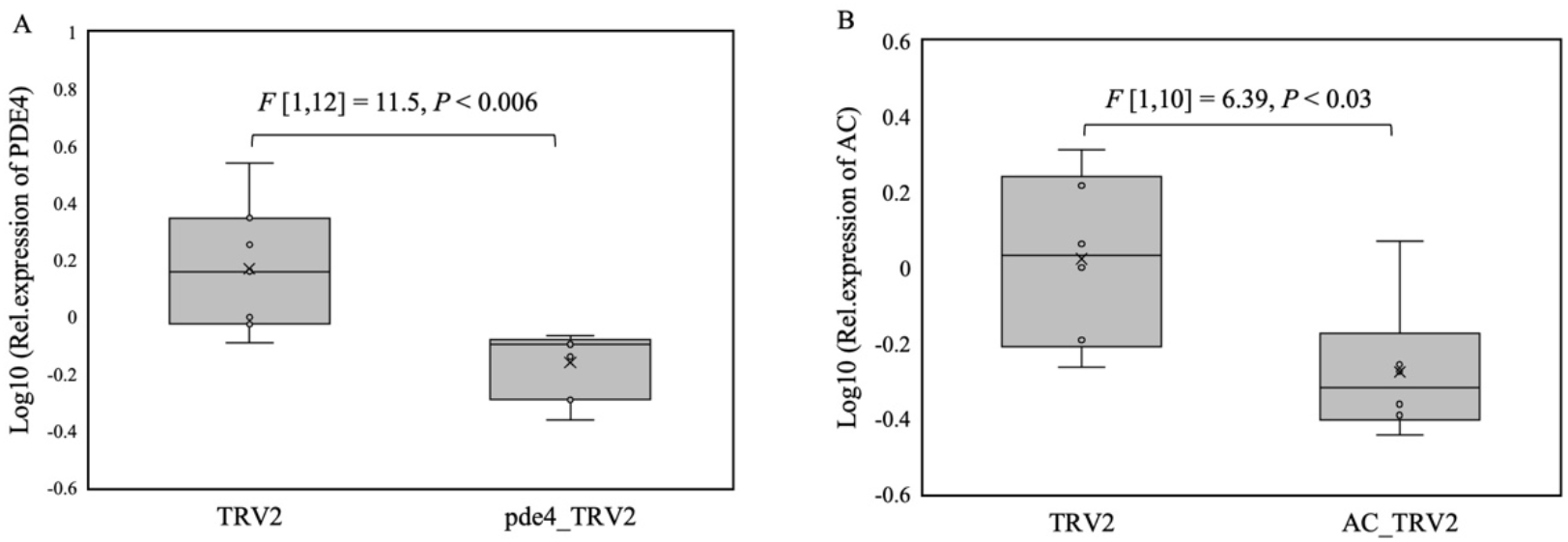
Expression of (A) PDE4 or (B) AC in F1 *B. tabaci* adults reared on tomato plants transformed with pde4_TRV2 or AC_TRV2 constructs compared with TRV2 control plants.

**Figure 8:**
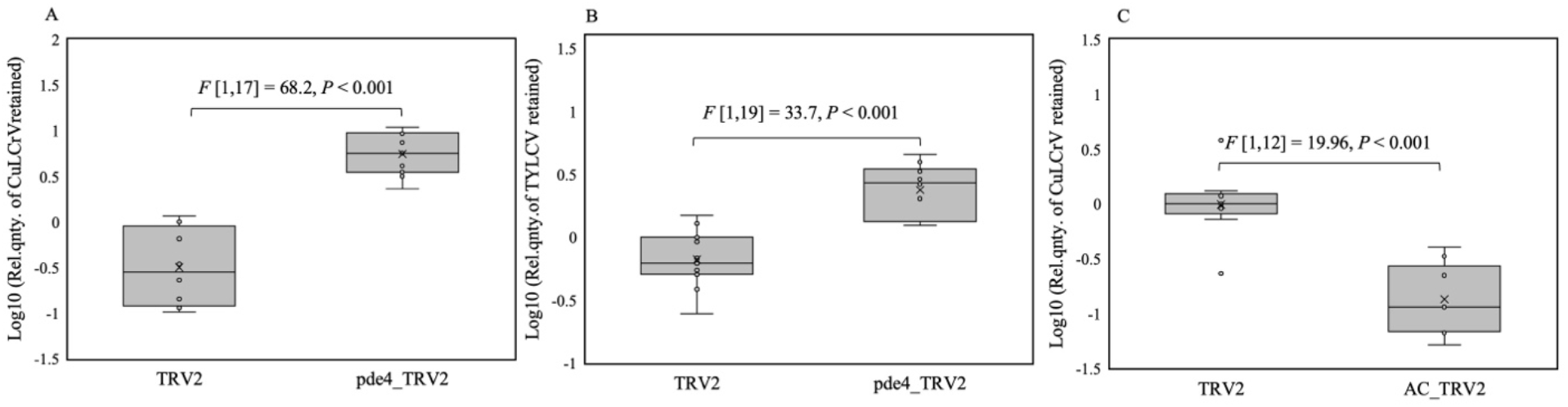
Relative quantities of (A) CuLCrV and (B) TYLCV retained (normalized to the □-tubulin gene of the whitefly) in F1 *B. tabaci* adults reared on either pde4_TRV2 or TRV2 plants after 24 hours of acquisition access to infected plants and overnight gut clearing on cotton plants. (C) Relative quantities of CuLCrV retained after 24 hours of acquisition and overnight gut clearing in F1 adults reared on AC_TRV2 compared to TRV2 control.

#### Transmission of CuLCrV is affected by cAMP levels within whitefly

To assess the effect of elevated cAMP on begomovirus transmission, whitefly adults were provided with an acquisition access for 48 hours on 20% sucrose diet with/without rolipram (200 □M) and provided with 24 hours of acquisition access on CuLCrV-infected plants were released to inoculate non-infected squash plants. Rolipram increased transmission efficiency (Fig. 9) of CuLCrV by *B. tabaci* adults in all three replications. Plants inoculated using rolipram-fed whiteflies (93.7%) differed significantly (□2 = 22.1, P<0.01) compared with plants infected (57.8%) using control whiteflies. Similarly, to evaluate the effect of reduced cAMP on begomovirus transmission, transmission efficiency of CuLCrV was compared between whitefly adults with acquisition access on a diet with and without SQ22536. Unlike rolipram treatment, no significant differences in transmission efficiencies between SQ22536 (22 infected out of 25 inoculated plants, 88%) or control diet (21 infected out of 24 inoculated plants, 87.5%) fed whitefly adults were observed. Further, CuLCrV loads in plants inoculated with rolipram/SQ22536 and control whiteflies were quantitated. Interestingly, virus loads in plants infected with rolipram-fed whitefly adults were significantly higher (Fig. 10A) than control. In contrast, plants infected with SQ22536-fed whitefly adults had significantly lower (Fig. 10B) virus loads compared with control.

**Figure 9:**
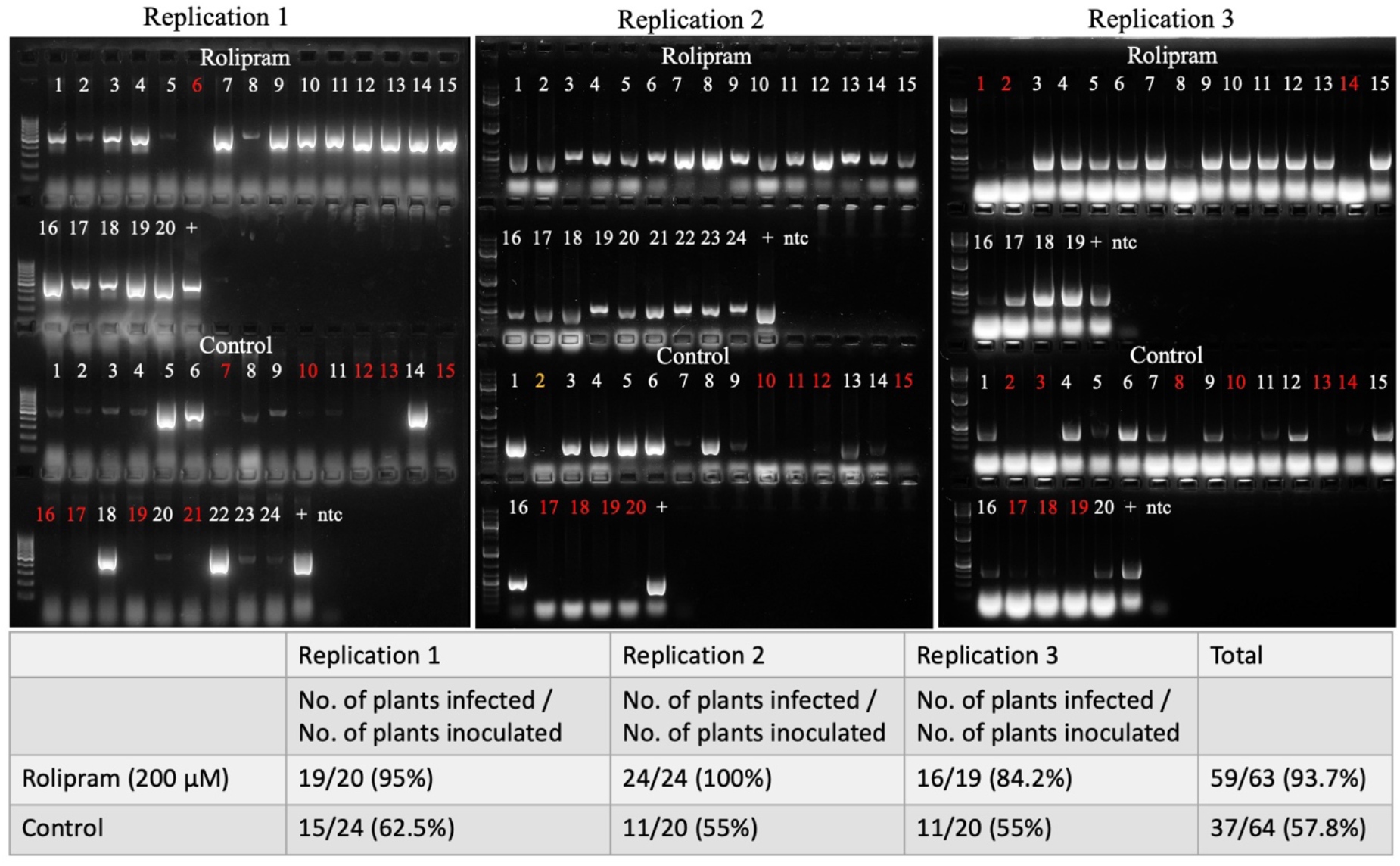
Transmission efficiency of CuLCrV by *B. tabaci* adults fed on rolipram or control diet and tested by PCR, 15 days post inoculation. The experiment was set up in three replicates for each treatment (rolipram/control diet) with a minimum of 19 plants inoculated in each replicate. The plant samples that tested positive for CuLCrV by PCR are marked in white numbers and the negative samples are marked in red. A positive (+) control DNA isolated from an CuLCrV and no template control (ntc) were used for the PCR test in each replicate.

**Figure 10:**
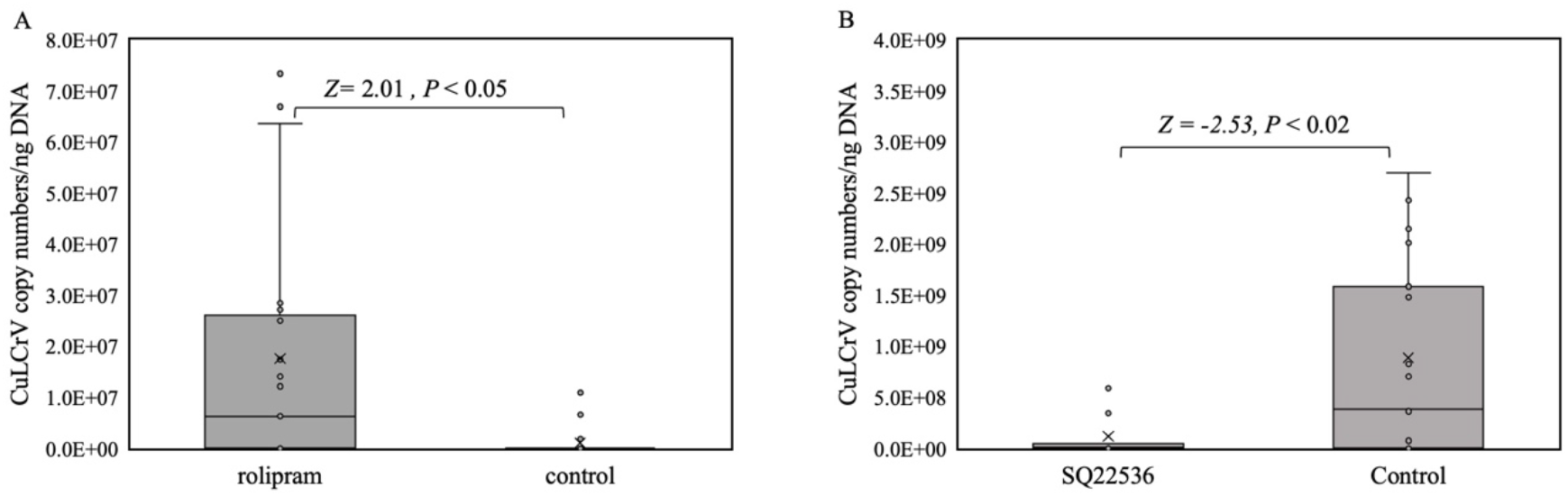
Quantitation and comparison of CuLCrV loads in infected squash plants inoculated with *B. tabaci* adults following an acquisition access (A) rolipram or (B) SQ22536 with that of whiteflies that fed on control diet.

## Discussion

Whitefly transmitted begomoviruses such as CuLCrV, SiGMV, and TYLCV cause severe economic losses to vegetable production in southeastern and southwestern United States. Disruption of genes/pathways critical for virus transmission can lead to effective management of begomoviruses. In this study, a gal4 based Y2H system was used to screen the cDNA library of *B. tabaci* (B cryptic species) with the CP of CuLCrV as a bait to identify interacting whitefly proteins involved in transmission of begomoviruses. This study identified PDE4, a cAMP specific 3’, 5’-cyclic nucleotide phosphodiesterase of the whitefly that interacts with the CPs of begomoviruses via Y2H, and confirmed the interaction through pull-down assay, co-immunoprecipitation, and co-immunolocalization in the whitefly midgut. PDE4 family of enzymes regulate intracellular cAMP, a second messenger regulating multiple downstream signaling cascades. Two enzymes, AC and cAMP dependent phosphodiesterase, maintain the intracellular cAMP concentration. Extracellular ligand activity with cell surface receptors such as the G-protein coupled receptor activates AC, which catalyzes the synthesis of intracellular cAMP from AMP, whereas PDE4 hydrolyzes cAMP to AMP (28). Intracellular cAMP concentrations are sensed by its binding with two intracellular receptors, Protein Kinase A (PKA) and Exchange Protein directly activated by cAMP (Epac). Cyclic AMP dependent activation or deactivation of these receptors orchestrate diverse metabolic and physiological functions of the cell. Immune response to infection with pathogenic microbes also can be modulated by intracellular cAMP concentrations (29). Elevated intracellular cAMP leads to downregulation of NF-kB induced transcription of inflammatory and apoptotic responses or upregulation of anti-inflammatory cytokines (30).

Pathogenic viruses (31,32) and bacteria (33) of humans are known to elevate cAMP levels within its host cell to evade immune response, and restoration of AMP leads to heightened immune response (34). Thus, intracellular cAMP has been an important target for therapeutic purposes for years (25). The results of this study demonstrate that elevation of cAMP in whitefly by inhibition of PDE4 causes increased begomovirus retention and transmission. Similarly, reduction in cAMP levels by inhibiting AC led to decreased begomovirus retention and lower virus loads in infected plants after transmission. Multiple transcriptomic studies have reported increased expression of immune related genes on acquisition of begomoviruses by the whitefly (35–38). Interactions of whitefly proteins with begomoviruses have been reported to either facilitate or inhibit virus transmission. For example, binding of CPs with heat shock protein 70 (HSP 70) (20) has been implied as a mechanism of the insect for degradation of virus particles, whereas binding to statin (39) has been implied as a virus strategy to prevent activation of JAK/STAT immune response. Interestingly, elevated cAMP is known to induce the expression of HSP 70 (40), whereas it inhibits JAK/STAT signaling pathway (41). However, results in this study do not explain the functional role of interaction of begomovirus CP proteins with the whitefly PDE4. Whether interactions between PDE4 and the CP is evolved to aid or to inhibit virus transmission remains elusive, as significant differences in cAMP levels between viruliferous and non-viruliferous whiteflies were detected. However, results of this study clearly show that elevated cAMP facilitates virus retention and transmission.

The findings of this study indicate potential for using insecticides/chemicals that reduce cAMP levels in the whitefly for management of begomoviruses. Bio-pesticides modulating intracellular cAMP of insects by activation of AC or inhibition of PDE have been used as insecticides previously (42). Further exploration and exploitation of such natural active ingredients can open new avenues for alternative and sustainable management of begomoviruses. Recent novel strategies, such as foliar application of nano-clay particles loaded with dsRNA targeting virus and insect genes have been shown to be effective for the management of plant viruses (43) and the whitefly vector (44). However, management of insect transmitted viruses by foliar dsRNA applications can be made more effective if the insect vector and virus transmission mechanism are simultaneously targeted. Aiming to decrease cAMP levels of the whitefly by targeting the AC gene through foliar application of RNAi insecticides can be an effective strategy to reduce begomovirus epidemics and will be investigated in the future.

## Author contributions

Conceptualization, S.G., R.S., and M.G.; methodology, S.G., B.M., S.Gm., O.J., R.S., and M.G.; validation, S.G., B.M.; formal analysis, S.G.; investigation, S.G., B.M., S.Gm., and O.J.; resources, R.S., and M. G.; writing—original draft preparation, S.G.; writing—review and editing, S.G., B.M., S.Gm., O.J., R.S., and M.G.; visualization, S.G.; supervision, R.S., and M.G.; project administration, R.S.; and funding acquisition, R.S. All authors have read and agreed to the published version of the manuscript.

## Funding

This work was supported by grants from the Georgia Commodity Commission for Vegetables, Georgia Department of Agriculture—Specialty Crop Block Grant, and the UGA-USDA ARS cooperative agreement number 6080-22000-027-18S.

## Notes

### Competing Interest Statement

The authors have declared no competing interest.

